# Amplicon and metagenomic analysis of MERS-CoV and the microbiome in patients with severe Middle East respiratory syndrome (MERS)

**DOI:** 10.1101/2020.11.28.400671

**Authors:** Waleed Aljabr, Muhannad Alruwaili, Rebekah Penrice-Randal, Abdulrahman Alrezaihi, Abbie Jasmine Harrison, Yan Ryan, Eleanor Bentley, Benjamin Jones, Bader Y. Alhatlani, Dayel AlShahrani, Zana Mahmood, Natasha J. Rickett, Bandar Alosaimi, Asif Naeem, Saad Alamri, Hadel Alsran, Maaweya Hamed, Xiaofeng Dong, Abdullah Assiri, Abdullah R. Alrasheed, Muaawia Hamza, Miles W. Carroll, Matthew Gemmell, Alistair Darby, I’ah Donovan-Banfield, James P. Stewart, David A. Matthews, Andrew D. Davidson, Julian A. Hiscox

## Abstract

Middle East Respiratory Syndrome coronavirus (MERS-CoV) is a zoonotic infection that emerged in the Middle East in 2012. Symptoms range from mild to severe and include both respiratory and gastrointestinal illnesses. The virus is mainly present in camel populations with occasional spill overs into humans. The severity of infection in humans is influenced by numerous factors and similar to severe acute respiratory syndrome coronavirus 2 (SARS-CoV-2) underlying health complications can play a major role. Currently, MERS-CoV and SARS-CoV-2 are co-incident in the Middle East and a rapid way is required of sequencing MERS-CoV to derive genotype information for molecular epidemiology. Additionally, complicating factors in MERS-CoV infections are co-infections that require clinical management. The ability to rapidly characterise these infections would be advantageous. To rapidly sequence MERS-CoV, we developed an amplicon-based approach coupled to Oxford Nanopore long read length sequencing. The advantage of this approach is that insertions and deletions can be identified – which are the major drivers of genotype change in coronaviruses. This and a metagenomic approach were evaluated on clinical samples from patients with MERS. The data illustrated that whole genome or near whole genome information on MERS-CoV could be rapidly obtained. This approach provided data on both consensus genomes and the presence of minor variants including deletion mutants. Whereas, the metagenomic analysis provided information of the background microbiome.

## Introduction

Coronaviruses were once described as the backwater of virology as they did not cause extensive disease in humans. However, with the emergence of severe acute respiratory syndrome coronavirus (SARS-CoV) in China in 2003, Middle East respiratory syndrome coronavirus (MERS-CoV) in Saudi Arabia in 2012 and now SARS-CoV-2 originating in 2019 in China, this is clearly not the case. These viruses cause respiratory and gastrointestinal illnesses and share similar genome architectures and disease profiles. Severe infection in humans is typified by cytokine storms^1,2^, pneumonia and kidney failure. MERS-CoV has been identified by the WHO as a prioritised disease and is on their list of pathogens for research and development in emergency contexts. With the advent of SARS-CoV-2 there is an urgent and unmet healthcare need to develop generic medical countermeasures to combat these infections and mitigate future outbreaks, particularly in contact tracing and understanding transmission dynamics. As a consequence, these viruses cripple national infrastructure, trigger city-wide transport curfews and disrupt international travel and commerce. Many countries are at risk for both SARS-CoV-2 and MERS-CoV.

There are striking parallels between the emergence of all three of these viruses. For example, one of the major concerns with MERS-CoV is the potential spread of the virus and other pathogens during Hajj^3^. This is an annual Islamic pilgrimage to Mecca in Saudi Arabia, involving some 2 million of the world’s population, and approximately 0.5 million Saudi residents. There are constant spill over events from camels into humans, and the geographical distribution of MERS-CoV in dromedaries is increasing either through spread and/or increased surveillance. Currently no antiviral therapies or vaccines are licensed for treatment or prevention of coronavirus infection in humans. Several studies are ongoing evaluating medical countermeasures for MERS-coronavirus infection. These have included compounds already in use in the clinic, such as combination of interferon-α2b and ribavirin^4^. A phase 1 DNA vaccine, based on the viral spike glycoprotein, funded by US Department of the Army, has been recently evaluated in a human trial^5^ and also in dromedary camels^6^. Additionally, a Phase 1 clinical trial has started in humans in Saudi Arabia assessing the safety and tolerability of a vaccine based on the ChAdOx1 MERS vaccine^7,8^. The sporadic nature of the outbreaks and the lack of suitable animal models has hindered research.

Given the lack of medical counter measures, shutting down transmission changes is essential in bringing such outbreaks under control^9^, and sequencing the genomes of viruses can aid in this through molecular epidemiology, as was demonstrated with SARS-CoV in Singapore^10,11^. This approach can also be used to identify properties of the virus relevant to health authorities, including the rate of virus evolution and whether targets of medical counter measures remain stable and valid. During the 2013-2016 West African Ebola virus outbreak, high resolution sequencing using Illumina^12^ and then long-read length/MinION^13^ based sequencing was used to provide measures of virus evolution^12^. The data also showed the origin of the outbreak was a single zoonotic transmission event and was spread through subsequent human to human transmission in Guinea and to surrounding countries^12,14^. The utility of the portable MinION based approach for rapid contact tracing was demonstrated in the West African outbreak through shutting down transmission chains arising from sexual contact^15^. Likewise, MinION based sequencing proved crucial in the 2018 Nigerian Lassa fever outbreak to show that independent transmission events from rodents rather than human to human contact was responsible for the spread of the outbreak^16^. MinION sequencing is also used for deriving consensus genomes for SARS-CoV-2 from clinical samples^17,18^.

Thus, providing sequence information of the genomes of viruses (or pathogens in general), is informative for health authorities, but only if the data is provided in real or near real time. Also, geopolitical considerations must be taken into consideration in terms of sample ownership and data sharing^19^, especially given the Helsinki Declaration. MERS-CoV is highly transmissible^20^ and sporadic outbreaks are ongoing in the Kingdom of Saudi Arabia and surrounding regions. Working with clinical specimens can prove challenging, particularly in understanding the pathology of MERS-CoV in how viral load and which co-infections influence outcome and the severity of this infectious disease. In order to aid in the diagnosis of MERS-CoV, and to rapidly generate viral genome sequence, the application of an amplicon-based and metagenomic MinION sequencing approach was to sequence viral RNA. During replication of coronaviruses the approximately 30 kb positive sense RNA genome, a nested set of subgenomic mRNAs are synthesized. Subgenomic mRNAs located toward the 3’ end of the genome are generally more abundant than transcripts nearer the 5’ end^21,22^. The data demonstrated that whole genome or near whole genome information on MERS-CoV could be rapidly obtained from samples taken from infected patients and sequenced using the amplicon approach. Further, the amplicon-based sequencing approach provided data on both consensus genomes and the presence of minor variants including deletion mutants. Metagenomic analysis provided information of the background microbiome and how this was associated with outcome. Real time analysis of the microbiome in patients and the identification of antibiotic resistance markers may provide better chemotherapeutic approaches to manage co-infections.

## Methods

### Ethics statement

Ethical approval was obtained from the Institutional Review Board no 18-102, King Fahad Medical City, Riyadh, Saudi Arabia. Informed consent to participant was not required since only remaining left-over samples were used for this study.

### Sample collection and processing

Nasopharyngeal aspirates (NPA), oropharyngeal, tracheal aspirates, throat swabs were collected from MERS-CoV positive patients (25–70 years old) admitted to different hospitals within Saudi Arabia. MERS-CoV diagnosis was confirmed by reverse transcriptase polymerase chain reaction (RT-PCR) (Biomerieux diagnostics). For this study, the NPAs with a confirmed MERS-CoV diagnosis from the Ministry of Health (MOH) Saudi Arabia, had no identifying information. The NPA, oropharyngeal, tracheal aspirates, throat swabs sampling were carried out as per the MOH’s guidelines. Samples were stored at −80°C until use. RNA from the NPA was extracted using EZ1 Virus Mini Kit v2 (Qiagen [955134]). The RNA concentration was measured by the Qubit RNA BA assay (Qiagen [32852]). Information on samples from patients use in this study is included in Table 1. This includes the sex, age, hospital, specimen type, Ct value of the E gene and ORF1AB, comorbidities, outcome and whether the patient was in an intensive care unit.

**Table 1:** Patient characteristics of patients included in this study, including sex, age, city, specimen type, Ct values for E-gene and RdRp, comorbidities, outcome and ICU status.

| Patient | Sex | Age | Hosp. | Specimen type | MERS-CoV Ct Value |  | Comorbidities | Outcome | ICU |
| --- | --- | --- | --- | --- | --- | --- | --- | --- | --- |
|  |  |  |  |  | E gene | Orf1 ab |  |  |  |
| 1 | Male | 64 | Medinah | Combined nasopharyngeal and oropharyngeal swab | 30.84 | 30.75 | Diabetes, Hypertension | Non-fatal | - |
| 2 | Male | 64 | Medinah | Combined nasopharyngeal and oropharyngeal swab | 33.75 | 33.47 | Diabetes, Hypertension | Non-fatal | - |
| 3 | Male | 80 | Medinah | Combined nasopharyngeal and oropharyngeal swab | 30.76 | 30.96 | - | Fatal | - |
| 4 | Male | 60 | Jeddah | Nasopharyngeal swab | 21.05 | 22.73 | Diabetes, Hypertension, Heart disease | Fatal | Yes |
| 5 | Male | 51 | Jeddah | Tracheal aspirate | 27.75 | 29.68 | - | Fatal | Yes |
| 6 | Male | 82 | Jeddah | Nasopharyngeal swab | 29.18 | 29.54 | Diabetes | Fatal | Yes |
| 7 | Female | 64 | Jeddah | Nasopharyngeal swab | 20.77 | 20.89 | - | Non-fatal | - |
| 8 | Male | 74 | Dammam | Combined nasopharyngeal and oropharyngeal swab | 23.49 | 24.81 | - | Fatal | Yes |
| 9 | Female | 74 | Dammam | Nasopharyngeal swab | 24 | 24 | Diabetes, Hypertension | Non-fatal | Yes |
| 10 | Male | 60 | Dammam | Combined nasopharyngeal and oropharyngeal swab | 19.11 | 21.26 | - | Fatal | Yes |
| 11 | Female | 45 | Medinah | Combined nasopharyngeal and oropharyngeal swab | 27.29 | 29.77 | - | - | - |
| 12 | Male | 61 | Medinah | Combined nasopharyngeal and oropharyngeal swab | 29.78 | 31.77 | Diabetes | Non-fatal | No |
| 13 | Male | 51 | Medinah | Combined nasopharyngeal and oropharyngeal swab | 29.27 | 31.78 | - | Non-fatal | No |
| 14 | Male | 60 | Medinah | Combined nasopharyngeal and oropharyngeal swab | 30.02 | 30.91 | - | Non-fatal | - |
| 15 | Male | 67 | Dammam | Throat swab | 30 | 31 | - | Fatal | Yes |
| 115 | Male | 46 | Riyadh | Nasopharyngeal swab | 23.37 | 23.98 | Diabetes, Hypertension | Non-fatal | Yes |

### Virus stock generation and infection

To prepare RNA from virus infected cells as a control for amplification, the MRC-5 cell lines was infected with MERS-CoV (ERASMUS strain) at a MOI of 5. Total RNA was purified using the Trizol method.

### DNase Treatment of RNA

RNA samples were treated with Turbo DNase (Invitrogen), by adding 1 μl of TURBO DNase with 5 μl 10x buffer in a 56 μl reaction. The reaction was incubated at 37°C for 30 minutes. 5 μl of inactivating agent was added and incubated at room temperature for 2 minutes. The reaction was centrifuged at 10,000g for 90s and the RNA supernatant was transferred into a new tube.

### Primer design and synthesis

Primers for the generation of overlapping amplicons were designed using the Primer3Plus platform and re-checked again by Primer Blast (NCBI) to avoid primers with a high self-complementary score. Primers were synthesized by Eurofins Genomics (the sequence of the primers is shown Table 2). The stock concentration was 100 μM, and primers were diluted in DNase/RNase free H_2_O to make a 10 μM working concentration.

**Table 2:**
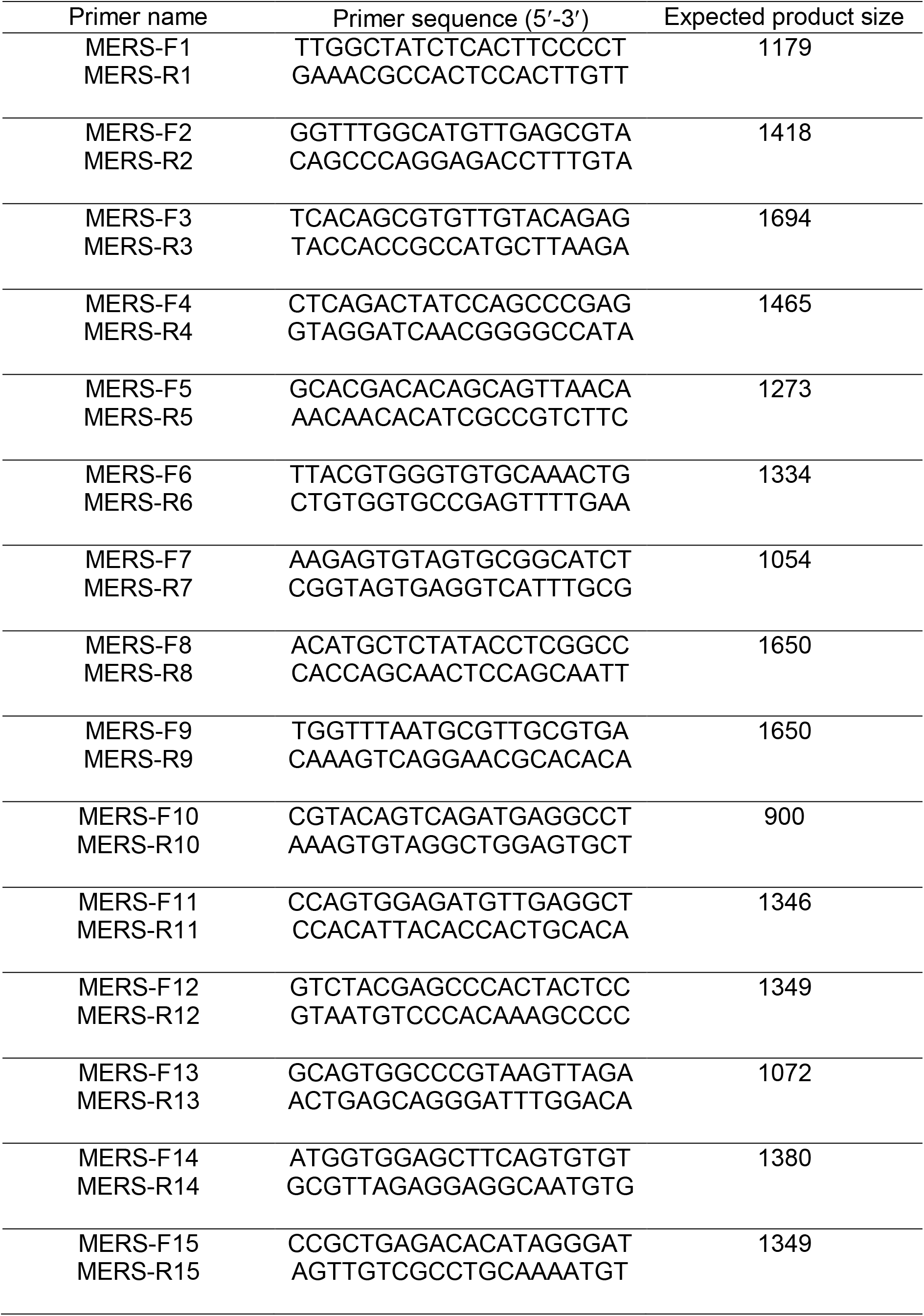
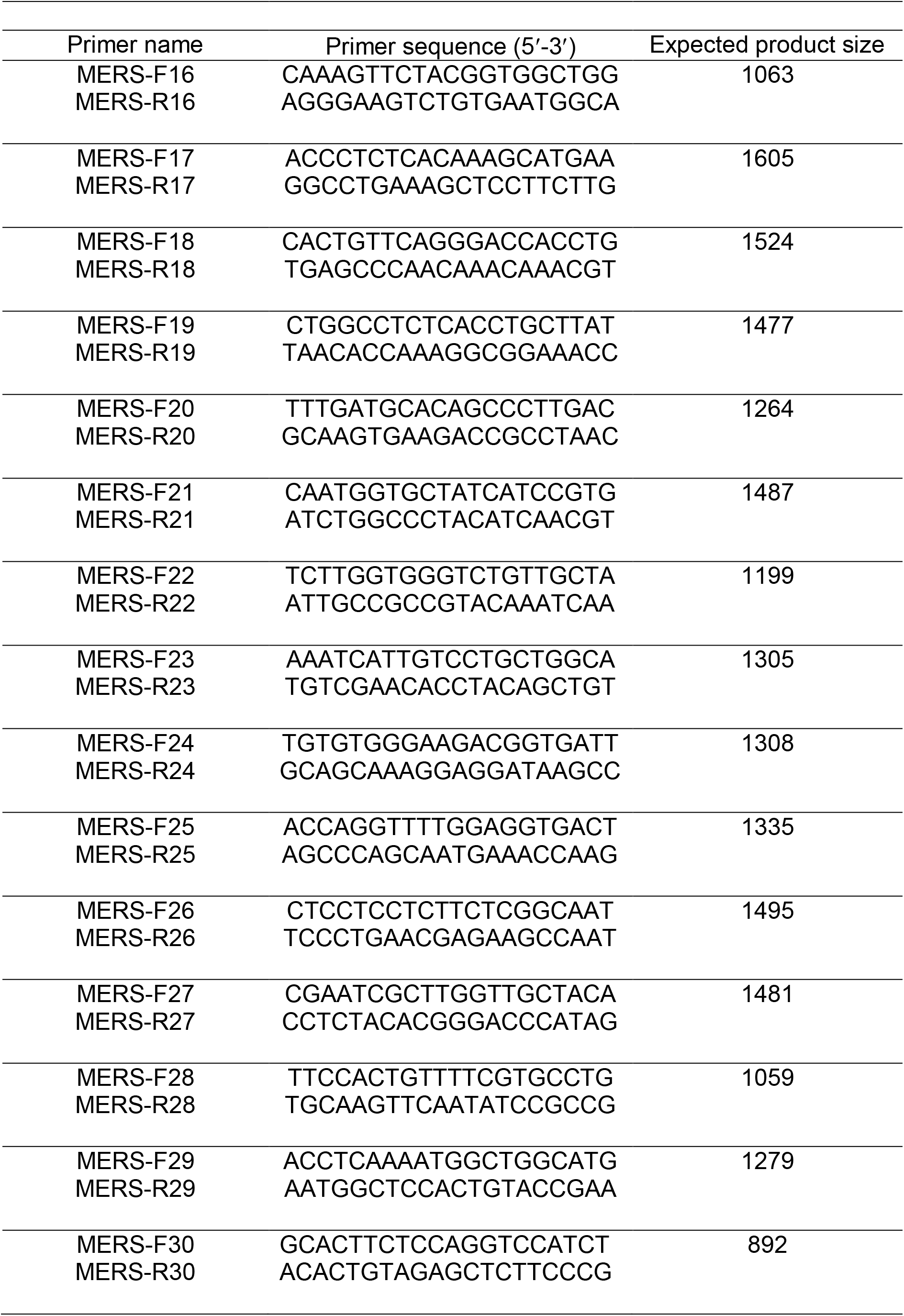
Details of primers sequence used in this study. The expected product size is given for the 30-amplicon approach (see Fig. 1).

**Fig. 1.**
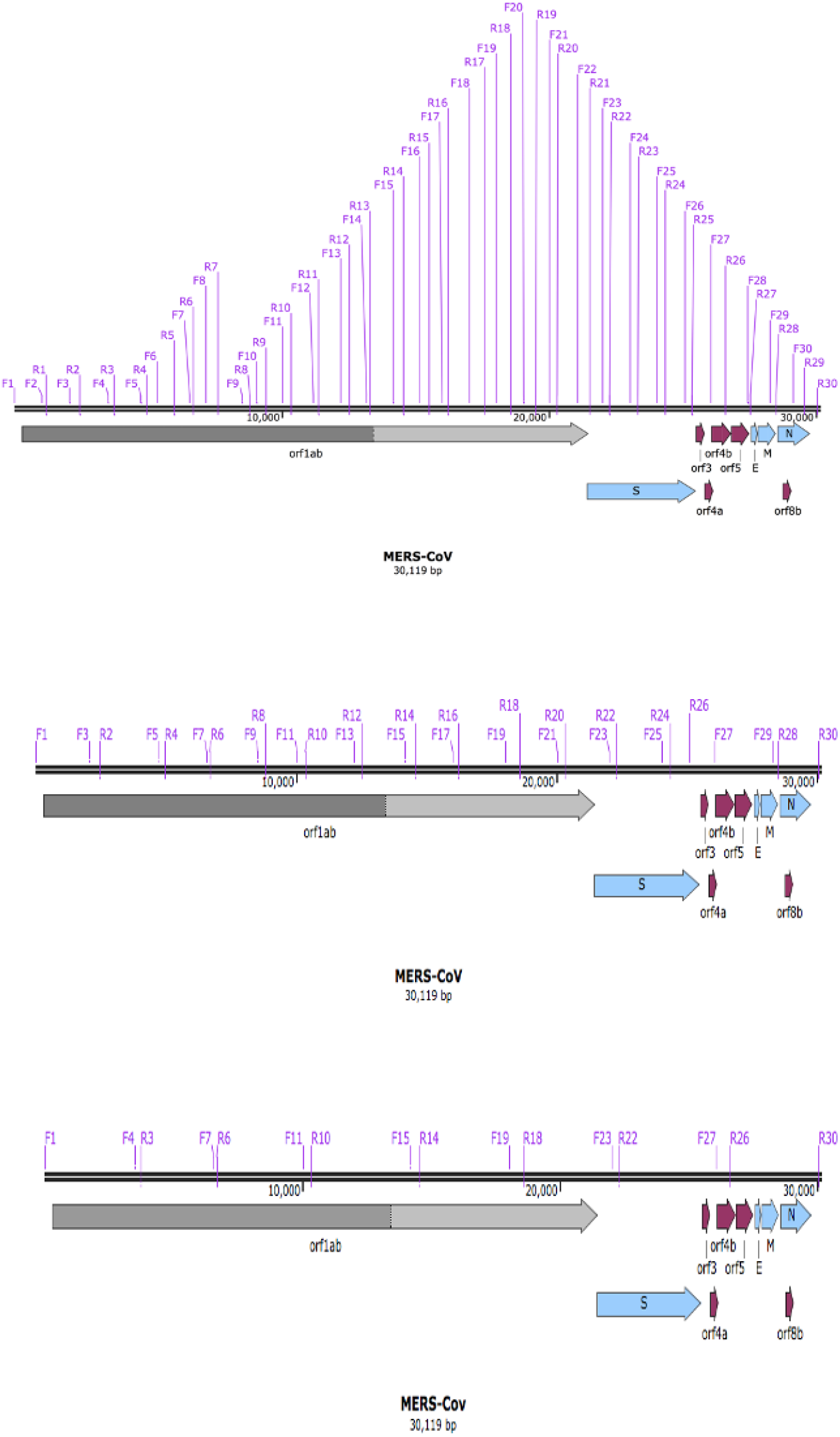
Location of conserved primer pairs (Table 1) on the MERS-CoV genome and position compared to MERS-CoV genes. Primer pairs can be used to generate amplicons of varying lengths including 30, 15 and 8 amplicons as indicated.

### cDNA synthesis and PCR

Superscript IV reverse transcriptase (Thermofisher [18090010]) and random hexamers (2.5 μM) were used to generate cDNA templates from RNA. cDNA was amplified using Q5-high fidelity DNA polymerase (NEB [M0491]).

### Amplicon analysis and sequencing

PCR products were run on a 1% agarose gel in TAE buffer to confirm the presence of amplicons. The amplicons generated from an individual patient were pooled and cleaned using AMPure XP Beads (Beckman Coulter [A63882]) following the manufacturer’s instructions before preparing the sequencing library with the Ligation Sequencing Kit (Oxford Nanopore Technologies [SQK-LSK109]). The sequencing library was added to a Flow cell connected to a MinIT and sequencing was initiated through MinKNOW. An initial rapid base calling setting was used.

### SISPA

RNA was reverse-transcribed with SuperScript IV Reverse Transcriptase (Invitrogen) using Sol-PrimerA (5′-GTTTCCCACTGGAGGATA-N9-3′), followed by second-strand DNA synthesis with Klenow (NEB). Reaction conditions for Round A were as follows: 1 μL of SolPrimerA (40 pmol/μL) was added to 4 μL of sample RNA, heated at 65 °C for 5 min, then cooled on ice for 5 min. Then 7 μL of SuperScript Master Mix (4ul 5XSSIV Buffer, 1ul 100mM DTT, 1ul RNaseOUT(Invitrogen) and 1ul SuperScriptTM IV Reverse Transcriptase (200U/ul) was added and incubated at 23 °C for 10 minutes, then 50°C for 10 minutes. Reaction was inactivated by incubating at 80°C for 10 minutes. For second strand synthesis, Klenow Fragment (NEB) as in Chrzastek et al. (2017) was used. The cDNA from each reaction was purified using a 1:1 ratio of AMPure XP beads (Beckman Coulter, USA). 5ml cDNA was amplified with Q5 High-Fidelity DNA Polymerase (NEB) according to manufacturer’s instructions with Primer Sol-B. Cycling conditions were as follows; 98 °C for 30 s, followed by 30 cycles of 98 °C for 10 s, 55 °C for 30 s, and 72 °C for 1 min, with a final extension at 72 °C for 10 min. All amplified cDNA was purified using a 1:1 ratio of AMPure XP beads (Beckman Coulter, USA), and resuspended in 25 ml. Purified cDNA was quantified using Qubit dsDNA HS assay (Invitrogen) according to the manufacturer’s instruction.

### Bioinformatics

Fast5 files were base called again using Guppy to mitigate for the lower accuracy associated with the rapid base calling setting initially used on the MinIT. To investigate deletions in the MERS-CoV genome, library reads were filtered by expected amplicon size and then aligned to the NCBI MERS reference NC_019843.3 using minimap2 as per the artic pipeline and SVIM was used to identify deletions^18^.

In addition, samtools was used to sort and index the alignment files, picard was used to remove amplification duplicates then a custom perl script was used count the nucleotides at each position on the reference genome, when the mapping quality was above 10. Data was visualised using R studio using the ggplot2. To consider the minor variation, the nucleotide depth at a single position with less than 20 counts was not taken forward into analysis to mitigate for random sequencing errors.

For rapid assessment of the meta-transcriptome approach, Fastq files were uploaded to Oxford Nanopore Technology’s (ONT’s) cloud-based pipeline EPI2ME (Fastq Antimicrobial Resistance; WIMP (rev. 3.4.0), ARMA CARD (rev. 1.1.6)) workflow to retrieve taxonomy classification and antimicrobial resistance information from MERS-CoV clinical samples.

In addition, meta-transcriptomic reads were assessed with Kraken2. Host removal was carried out to remove human reads. The Human genome assembly GRCh38.p13 was used as a reference. The Human genome assembly was indexed with minimap2 using the parameter “-ax map-ont”^23^. The quality-controlled Oxford Nanopore fastq reads were mapped to the indexed human genome assembly with minimap2 using default parameters. The resulting sam file was sorted and converted to bam with the command “samtools sort”^24^. A fastq file with unmapped reads was produced with the “samtools fastq” command with the parameter “-f 4”^24^

The quality controlled, host removed reads were classified using Kraken2^25^. The standard kraken output and sample report output files were produced with the options “--output” and “--report” respectively. The database used during Kraken classification consisted of the “bacteria” and “viral” kraken2 reference libraries (built 23rd April 2020). Visualisation of the Kraken2 output was carried out with Pavian, utilising the Kraken2 report files^26^. Kraken-biom (https://github.com/smdabdoub/kraken-biom) was used to generate a biom table which was then read into R studio with Phyloseq to plot alpha diversity and abundance of species identified in the clinical samples at the genus level^27^. DESeq2 was used to compare the difference in abundance of species between fatal (n=7) and non-fatal (n=7) MERS-CoV infections^28^.

## Results and Discussion

Coronaviruses encode several proteins that are involved in proof reading during virus replication^29,30^, therefore using nucleotide divergence for contact tracing may be problematic. Recombination and resulting deletions (and potentially insertions) may account for the wide genome diversity observed in some strains of coronaviruses^31-33^. These recombination events can led to new strains of coronaviruses^34^ or potentially affect vaccine strategies^35,36^. In order to identify these genome changes with sequencing, rapid long read length approaches offered by Oxford Nanopore were advantageous. Sample quality can also inform design of sequencing strategies and a RT-PCR approach was used to recover viral genome/transcript information. Rather than designing very short amplicons, longer amplicons were selected through selection of appropriate primer pairs along the MERS-CoV genome (Fig. 1).

### Primer design

Twenty MERS-CoV genome sequences were aligned with a reference sequence (NC_019843.3 – the ‘Erasmus Medical Centre (EMC)’ sequence). These sequences represented viruses collected in regions of Saudi Arabia and countries that reported cases including South Korea. Primer binding sites were chosen from conserved regions after alignment, so that a minimum of roughly 1000 bp sequential amplicons would be generated with an approximately 200 bp overlapping region at each terminus. This resulted in the selection of thirty sets of primer pairs (Table 2) that could be used to walk across the MERS-CoV genome with overlap between each generated amplicon. Alternatively, primer pairs can be selected that allowed the generation of larger amplicons that spanned more of the genome in contiguous reads (Fig. 1).

### Validation of primers and generation of amplicons using total RNA purified from MERS-CoV infected cells

To evaluate the utility of the selected primers for the amplification of viral RNA under controlled conditions, RNA was purified from MRC-5 cells that had been infected with the EMC strain of MERS-CoV at a MOI of 5. Infection was carried out under CL3+ conditions and total RNA purified from these infected MRC5-cells at 16 hrs post-infection. This RNA was used as a template to prime cDNA synthesis using random hexamers. Primer pairs (Table 2) were tested by gradient PCR to determine the optimum annealing temperature (data not shown). This resulted in amplification conditions (Table 3) such that the MERS-CoV genome was amplified using either 30 amplicons (Fig. 2A), 15 amplicons (Fig. 2B) or 8 amplicons (Fig. 2C), using the same set of conditions for each set of amplicons (Table 2). The rationale being that using the same amplification conditions across all primer pairs would be more efficient if large scale sample analysis was required. The data indicated that for the 30-amplicon approach, PCR products were observed that spanned the MERS-CoV genome. For the 15 and 8 amplicon approach products were also observed that spanned the MERS-CoV genome. However, amplification of these products varied in efficiency.

**Table 3:** PCR condition for 30, 15, and 8 amplicon approach.

| Step | Temperature | Time | Note |
| --- | --- | --- | --- |
| Initial denaturation | 98C° | 30 sec |  |
| Denaturation | 98C° | 10 sec | For 30*, 15**, or<br>8*** amplicon<br>approach |
| Annealing | 66C° | 30 sec |  |
| Extension | 72C° | 50*, 90**, or |  |
| 35 cycles |  | 180*** sec |  |
| Final extension | 72C° | 2 min |  |

**Fig. 2.**
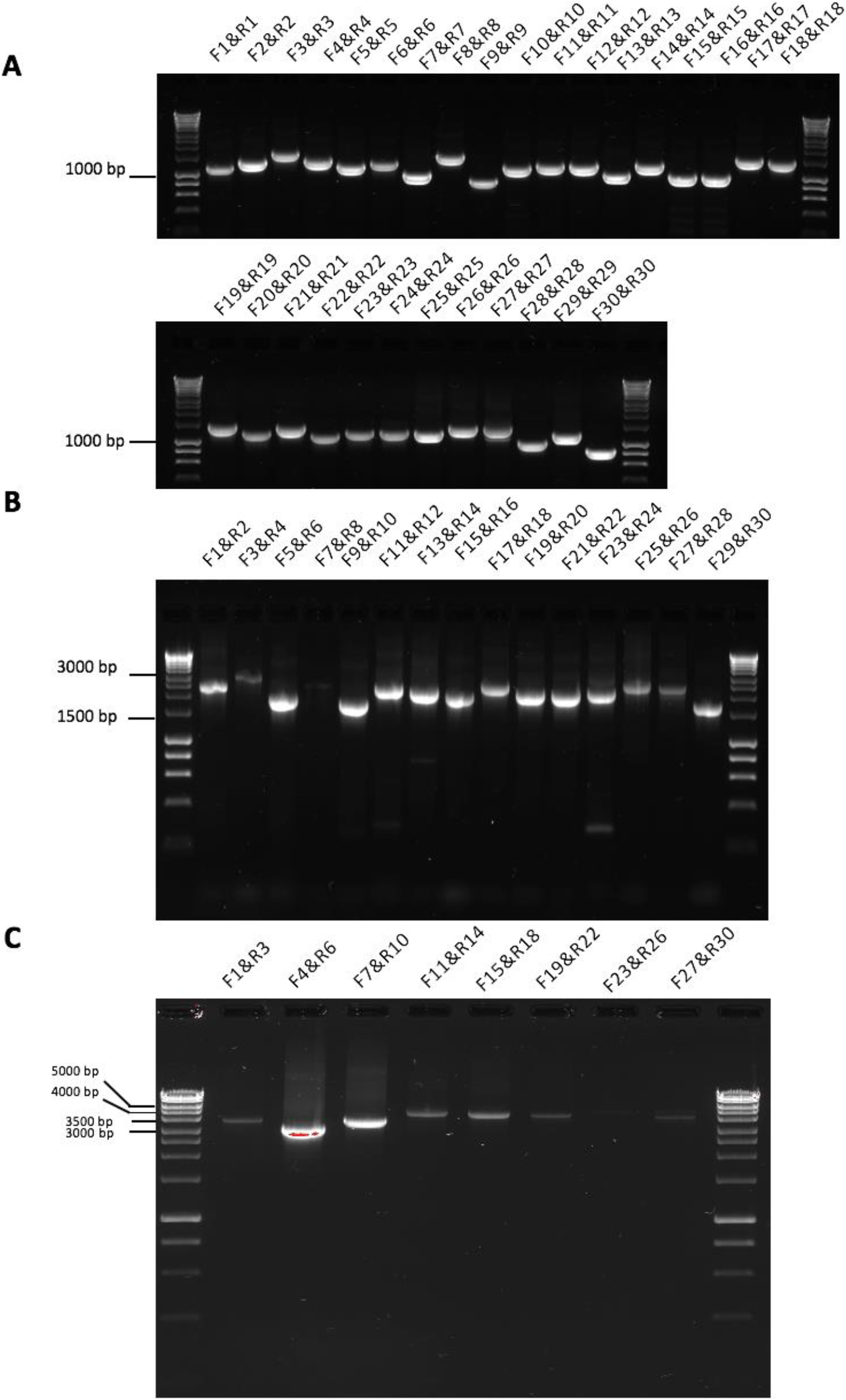
Agarose gel electrophoresis of amplicons generated using 30 (A), 15 (B) and 8 (C) combinations of primers pairs. These primer pairs were used to generate amplicons in combination with reverse transcription of RNA extracted from MERS-CoV infected cells.

### Generation of amplicons from patients infected with MERS-CoV and derivation of consensus genome sequence

The 30, 15 and 8 amplicon approach was evaluated to generate PCR products from MERS-CoV RNA extracted from nasal aspirates taken from patients with MERS. The rationale for MERS-CoV genome coverage using different size amplicons was to provide maximum versatility for walking across the genome to obtain sequence information. Note that this approach would also use the nested set of subgenomic mRNAs to capture sequence information. The data indicated that the 30 and 15 approach could be used to generate fragments from clinical samples (Fig. 3A and 3B, respectively). The 8-amplicon approach was not sufficient for obtaining coverage across the MERS-CoV from a clinical sample (data not shown). The PCR products generated in the 30 and 15 amplicon approach (from separate patients) were combined for each patient, barcoded and sequenced on separate flow cells for each patient. Sequencing reads generated by the MinION were aligned to a reference sequence. The analysis showed that complete genome sequence could be obtained from the 30 amplicon (Fig. 4A) and 15 amplicon approach (Fig. 4B). Consensus sequence was generated using the ARTIC bioinformatic pipeline. A custom perl script was used to count the number of each nucleotide against the reference sequence (NC_019843.3), to identify the viral genome sequence that dominated the population. This sequence was used to generate a bespoke consensus genome for an individual patient.

**Fig. 3.**
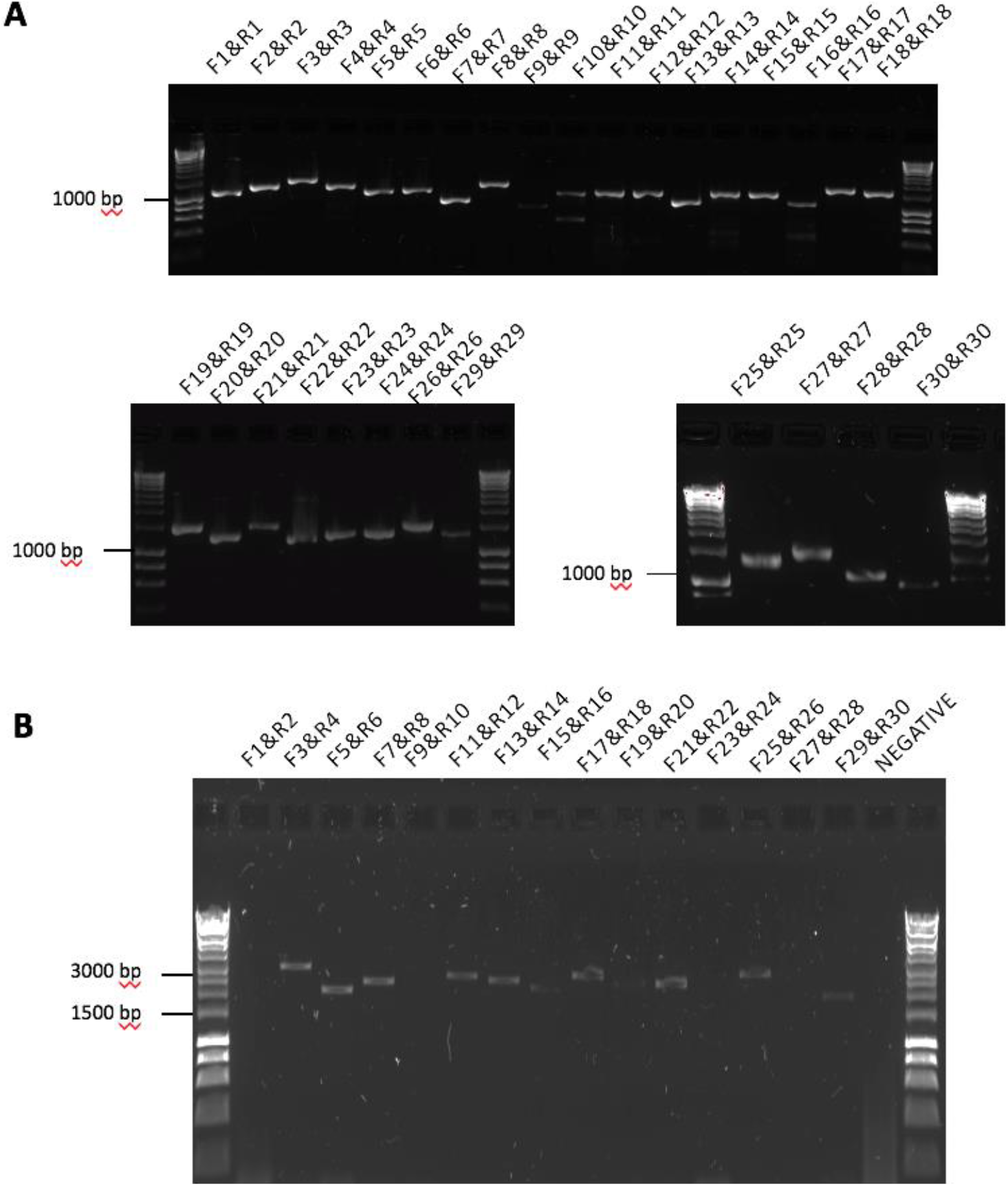
Agarose gel electrophoresis of amplicons generated using 30 (A) and 15 (B) combinations of primers pairs. These primer pairs were used to generate amplicons in combination with reverse transcription of RNA extracted from nasal aspirates taken from patients with MERS.

**Fig. 4.**
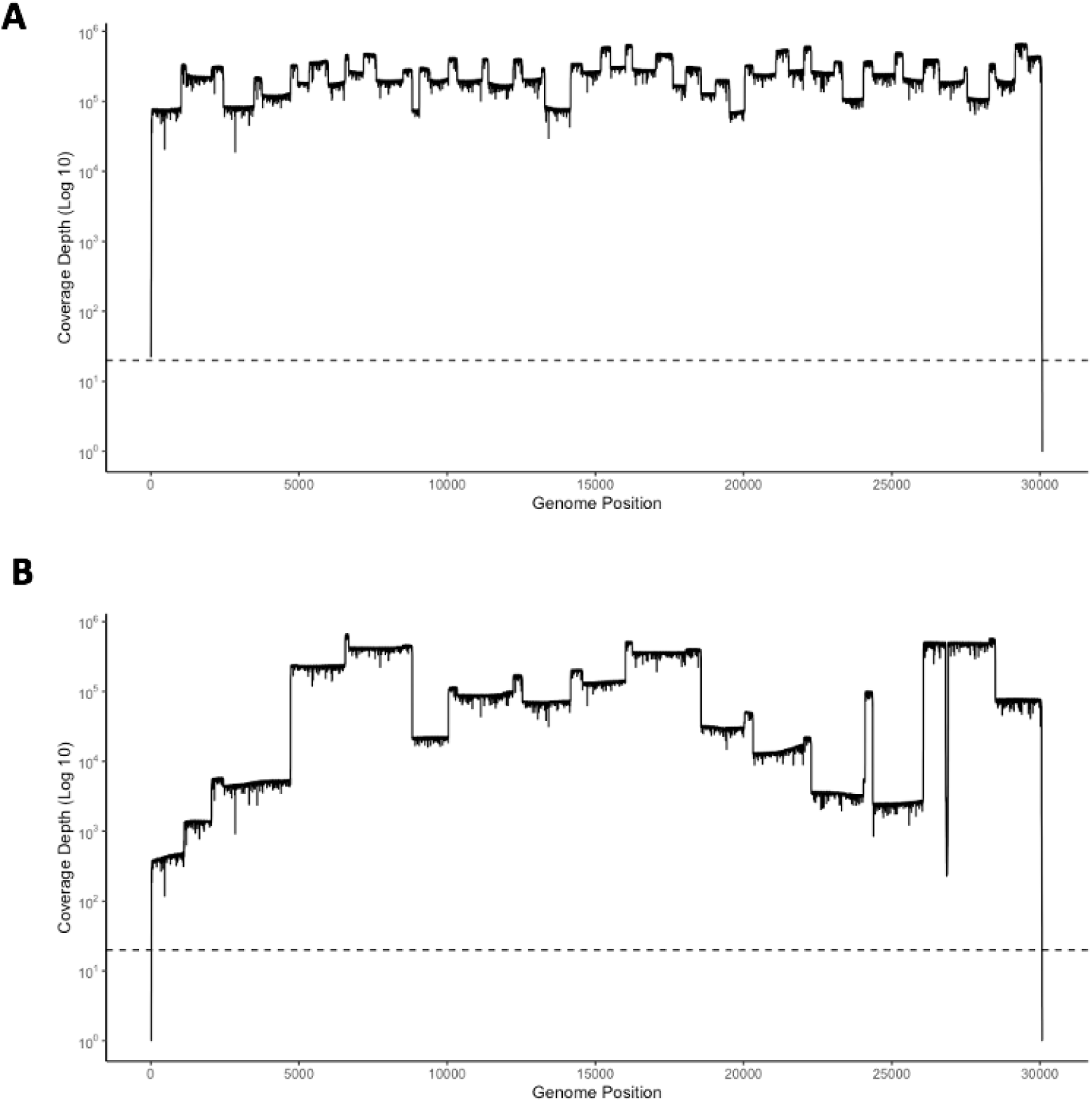
Read depth analysis of 30 (A) and 15 (B) amplicons sequenced on a single flow cell using Oxford Nanopore. Coverage of each position on the MERS-CoV genome is indicated on the y axis. Dashed line represents 20X coverage, indicating that above this line each nucleotide was sequenced at least twenty times.

### Analysis of the minor variant population within patients

The nucleotide substitution rate can drive the selection of genotypic and phenotypic variants of MERS-CoV. Whilst variation and potential functional changes in consensus genomes have been compared between patients, the ability to monitor minor variants and their frequency and how these contribute to the overall viral phenotype within a patient is unknown. The minor variant population in infection has been shown to influence the kinetics of virus replication and be associated with patient outcome^37^. Therefore, methodologies were developed that could be used to assess the minor variant frequency within a sample from a patient. The custom perl script used to call the consensus also revealed the nucleotide depth and the counts of each nucleotide at each position (Fig. 5). The depth was used to normalise the mutation frequency into a proportion instead of a raw count, allowing comparison for samples of different read depth. Nucleotides that had a count less than 20 were removed from analysis. As proof of principle, this approach was applied to the sequencing data obtained from patients 10 and 115. Patient 10 appeared to have more base changes in comparison to patient 115 (Fig. 6). Transitions (A>G, G>A, C>U, U>C) were more frequently observed and C>U seemed more prominent than other mutations. This is consistent with mRNA editing by APOBEC. APOBEC3 family members have been shown to be involved in restricting the growth of human coronavirus NL63 (HCoV-NL63)^38^ and identified as potential drivers of C to U transitions in SARS-CoV-2^39^. We would note that assessing minor variant populations using data from MinION sequencing may be problematic due to the higher error rate in sequencing than for example Illumina based approaches^13^. However, with sufficient read depth in a sample, an overview of the minor variant frequency/population can still be obtained.

**Fig. 5.**
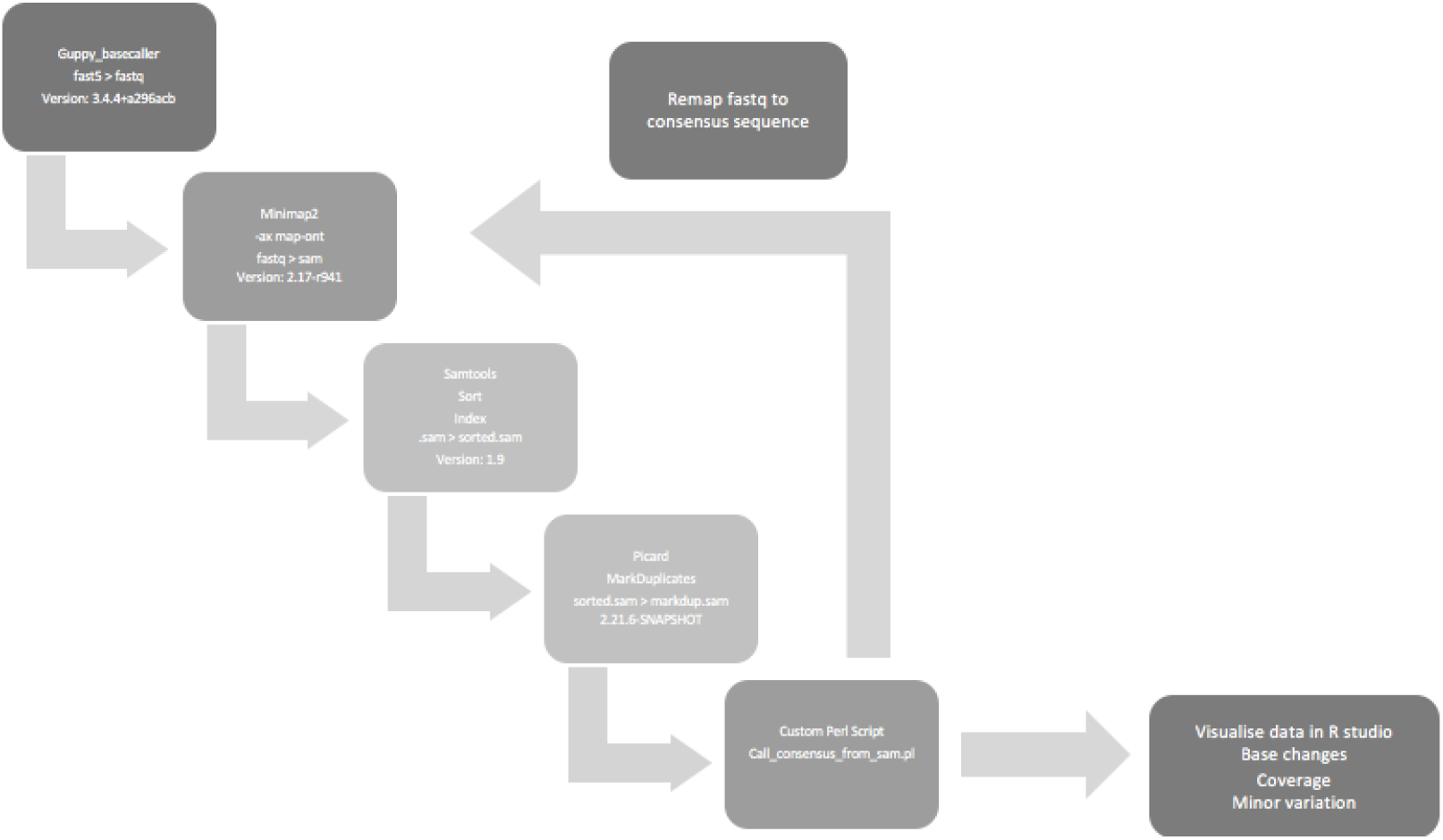
A flow diagram of the bioinformatic pipeline that was used to derive viral genome coverage and minor variation information of viral genomes within patients.

**Fig. 6.**
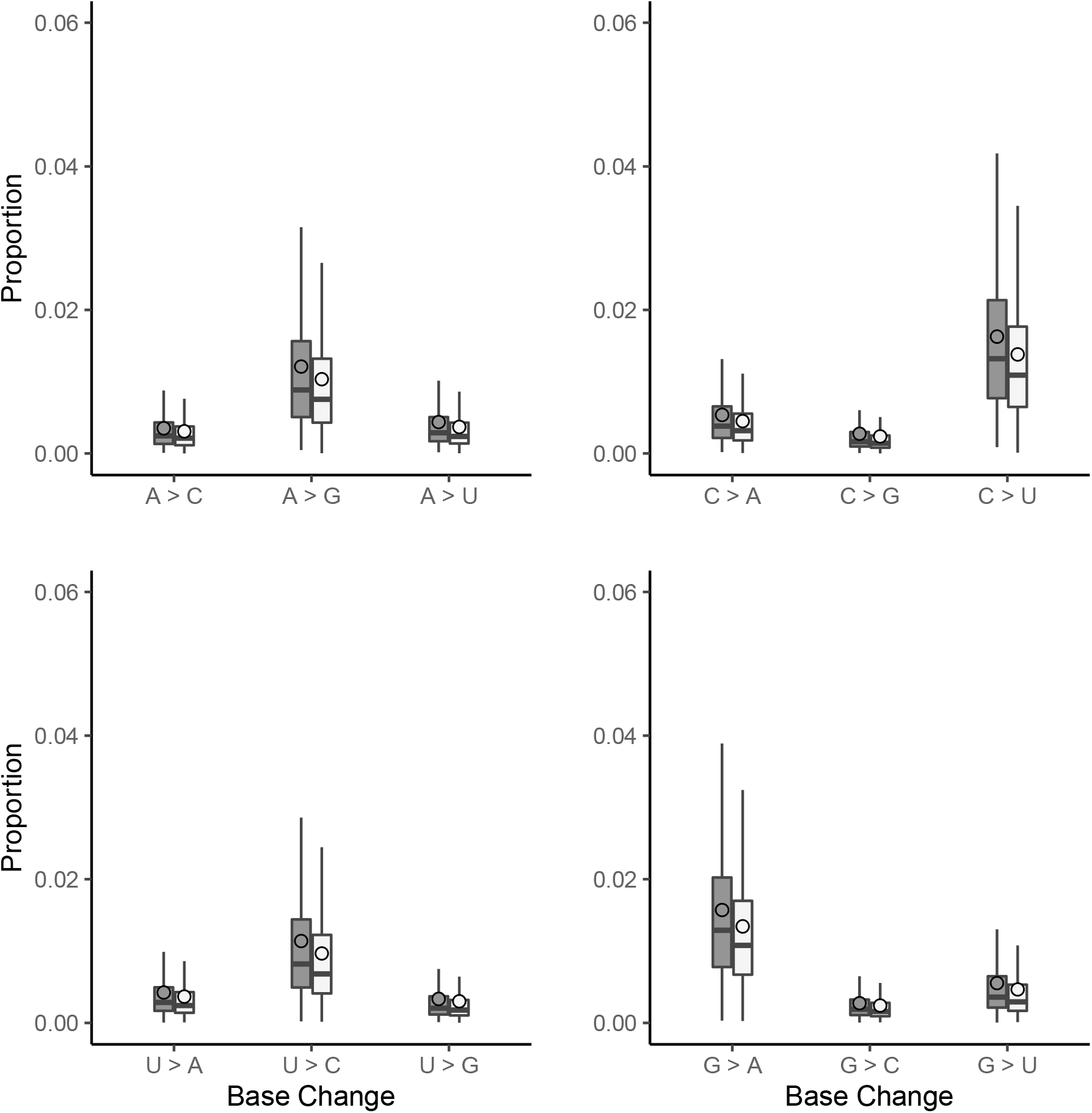
The sequencing reads were mapped to the patient consensus viral genome sequence. The custom script counted the number of each base at each genome position with a quality score >10. Positions with a depth <20 were removed from the analysis. This figure shows the proportion of base changes observed in comparison to the patient’s dominant consensus reference genome. Overall, transitions were observed more frequently than transversions, where C>U is the most observed base-change. We hypothesise that although transitions are more common, that APOBEC may have an influence on the MERS-CoV genome. Patient 10; dark grey, Patient 115; light grey, outliers not visualised.

**Fig. 7:**
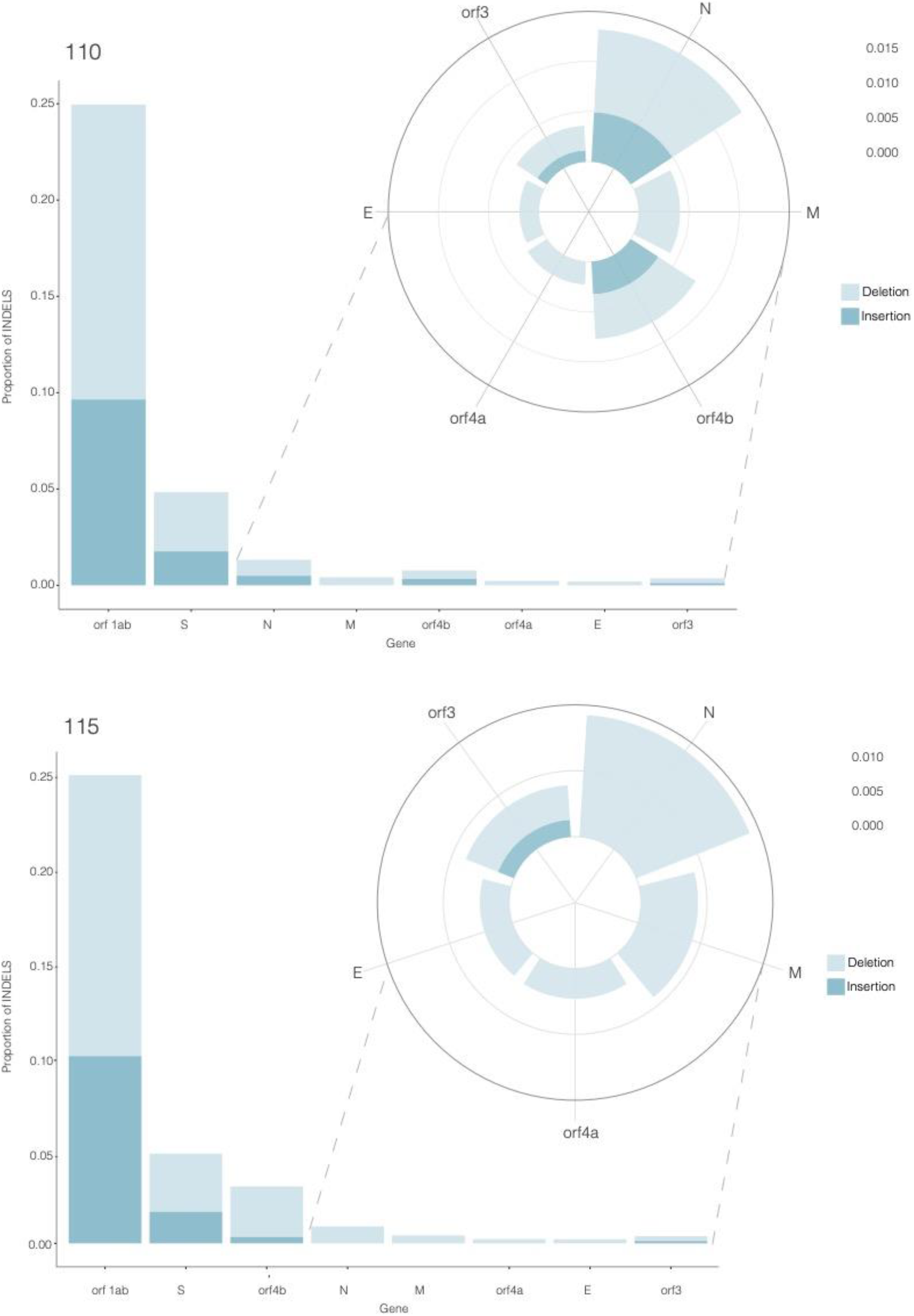
Deletions were assessed at the minor variant level as proportions in addition to the analysis in Table 4.

**Fig. 8:**
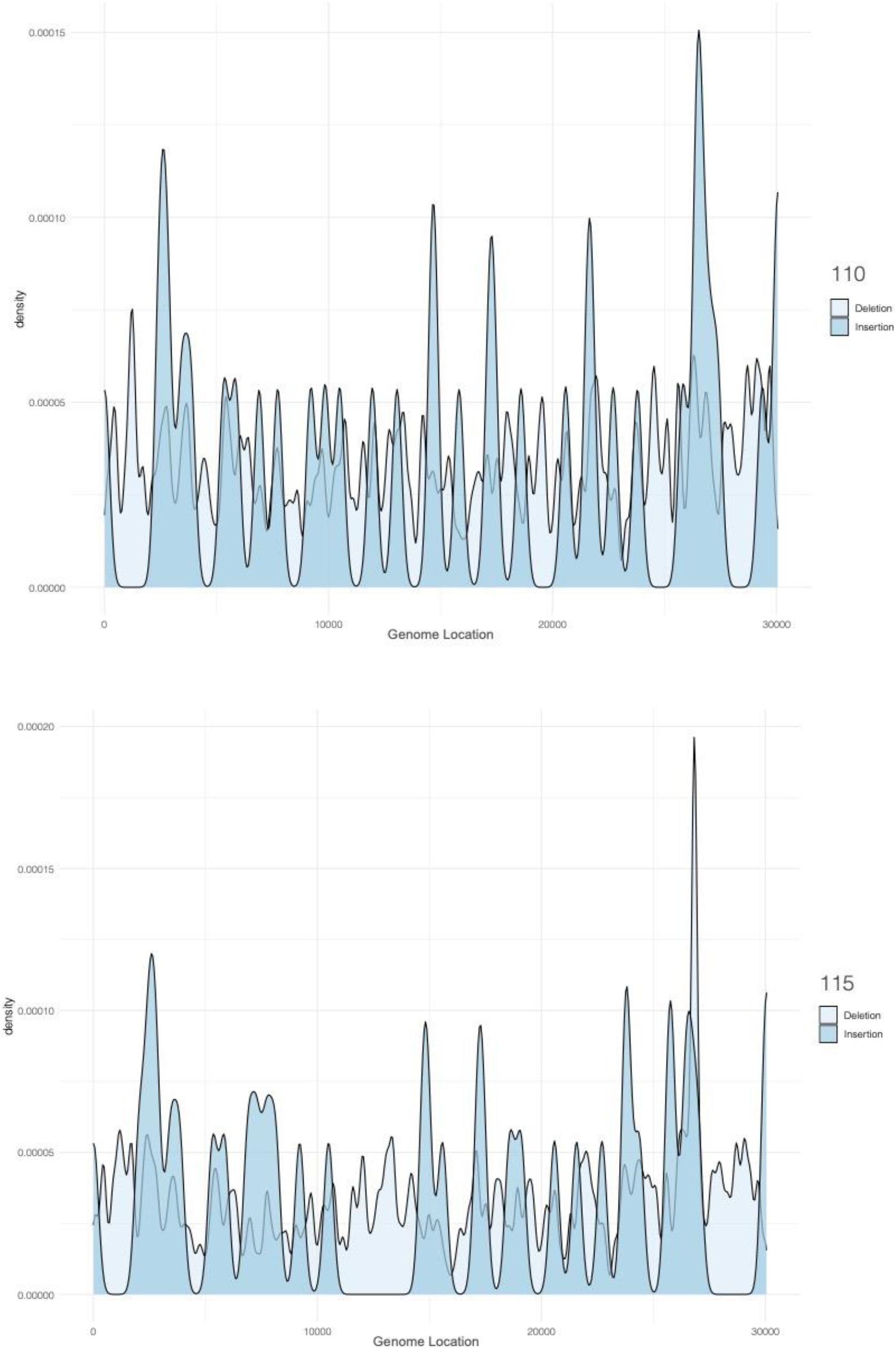
The density of indels at the minor variation level were plotted against genome position using sequencing data from patients 10 and 115.

### Identification and analysis of deletions in the viral genome in samples from patients

Sequencing data from patients 10 and 115 were interrogated for deletions using SVIM. Table 4 shows deletions identified with more than 5 supporting reads. Patient 10 had five deletions in Orf1ab and a deletion spanning N. Patient 115 was sequenced with the 15-amplicon approach and therefore generated amplicons over 2kb in length. A deletion of 77 bases that spans the orf4b and orf5 gene was identified in patient 115. Deletions in this region may have implications on virus pathogenesis, ORF4A is able to inhibit early antiviral responses (IFN α/β) in the host^40^, likewise ORF5 has been characterised to reduce inflammation responses^41^. This finding was consistent in both bioinformatic pipelines that were used. Several deletion candidates were identified in patient 10 in Orf1ab and N.

**Table 4:** Analysis of deletions present in the MERS-CoV genome from patients 115 and 10. Columns from left to right; Patient number, deletion start position (bp), deletion end position (bp), the number of supporting reads for this deletion, the quality score (which takes into consideration the mapping quality scores, where a value greater than 10 has higher confidence), standard deviation (SD) of the deletion span (bp) and SD of the position of the deletion from the supporting reads. Coordinates are given for the affected gene, and in the case of overlap, the second gene is provided.

| Deletion information |  |  |  |  |  |  | Affected Gene information |  |  |  |  |  |  |  |
| --- | --- | --- | --- | --- | --- | --- | --- | --- | --- | --- | --- | --- | --- | --- |
| Patient | Start (bp) | End (bp) | Supporting Reads | Quality Score | SD span | SD pos | Gene Start | Gene End | Gene Name | Overlap (bp) | Gene Start | Gene End | Gene Name | Overlap (bp) |
| 115 | 26820 | 26897 | 97 | 99 | 1.96 | 1.83 | 26093 | 26833 | Orf4b | 13 | 26840 | 27514 | Orf5 | 57 |
| 10 | 1315 | 2149 | 6 | 7 | 0.41 | 0.2 | 279 | 21514 | Orf1ab | 834 |  |  |  |  |
|  | 3576 | 4551 | 10 | 12 | 3.46 | 57.21 | 279 | 21514 | Orf1ab | 975 |  |  |  |  |
|  | 7611 | 8507 | 31 | 38 | 0.62 | 0.23 | 279 | 21514 | Orf1ab | 896 |  |  |  |  |
|  | 8818 | 9056 | 22 | 27 | 2.63 | 3.02 | 279 | 21514 | Orf1ab | 238 |  |  |  |  |
|  | 9407 | 10044 | 14 | 17 | 0.27 | 0.4 | 279 | 21514 | Orf1ab | 637 |  |  |  |  |
|  | 28514 | 29110 | 40 | 49 | 56.94 | 15.59 | 28566 | 29807 | N | 544 |  |  |  |  |

### Identification of bacterial and viral sequences in samples from patients and antibiotic resistance markers

RNA was extracted from respiratory samples taken from 15 patients with severe MERS, including combined nasal and oropharyngeal swabs (n=9), throat swab (n=1), nasal swabs (n=4), and tracheal aspirate (n=1). Metagenomic approaches have been used to identify the microbiome in samples from patients and correlate these with patient outcome^42^. SISPA was used to identify bacterial and viral transcripts present within the clinical samples. This was assessed using the real-time analysis pipelines Epi2Me with AMR or Kraken and Phyloseq packages. Species such as *Acinetobacter baumanni, Pseudomonas aeruginosa, Streptococcus pneumoniae* were identified, all of which can be associated with bacterial pneumonia. Fig. 9 illustrates the top 20 species identified across all 15 samples. Although detection of RNA does not always infer active infection, it provides insight of the potential causative agent for a bacterial pneumonia if one was to emerge.

**Fig. 9:**
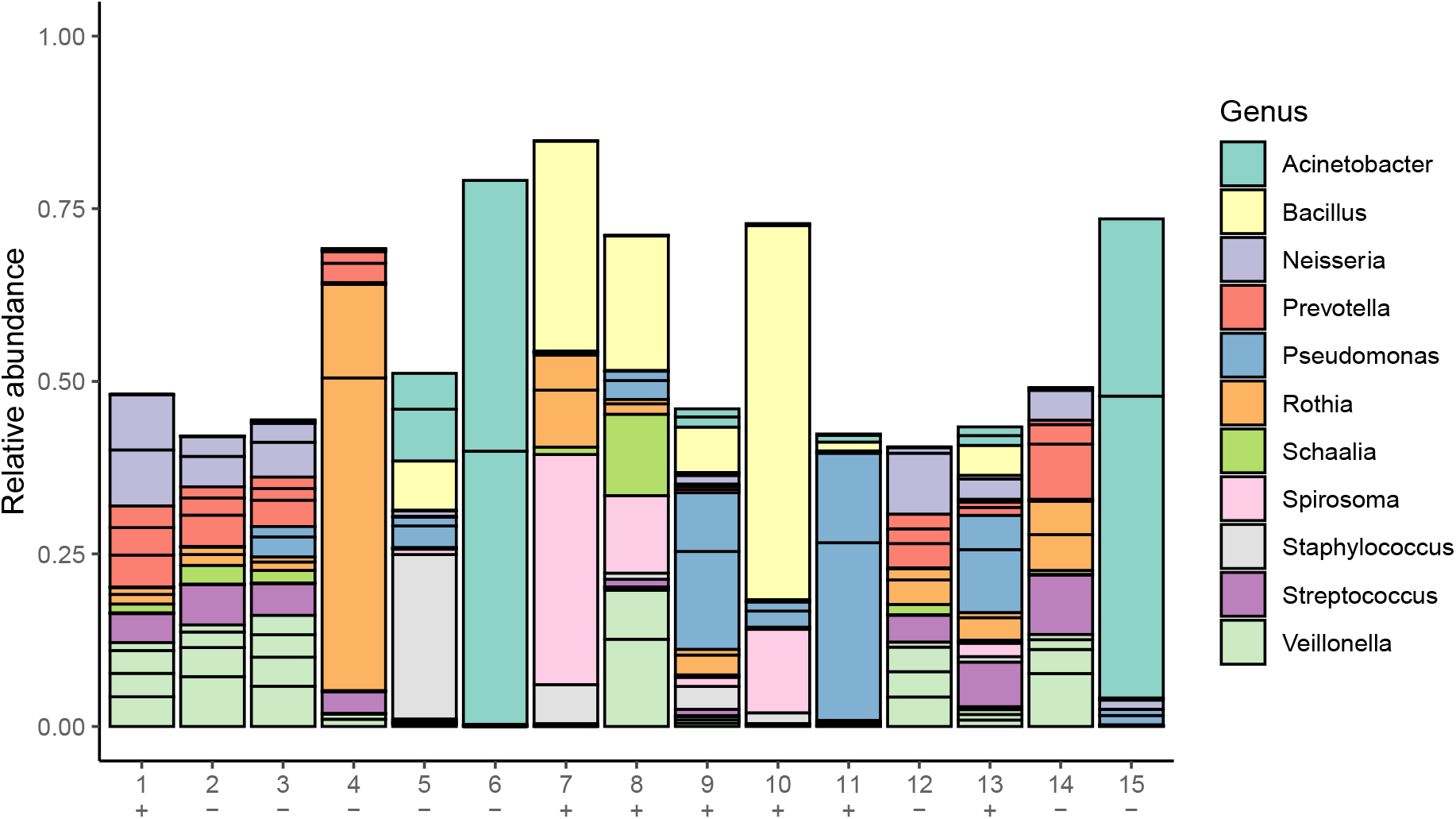
The top 20 species categorised into genus from 15 patients with severe MERS infection. Human reads were removed from sequencing libraries, and viral and bacterial transcripts were identified using Kraken2. Kraken2 outputs were converted into biom format before importing into R with Phyloseq. Relative abundance of each species is plotted for each patient. (+) MERS-CoV reads detected, (−) MERS-CoV reads not detected.

Patients with *Acinetobacter* transcripts identified were associated with fatal outcome of disease (Fig. 9). Acinetobacter infections are often exclusive to healthcare settings, especially in patients who have received ventilation support. Acinetobacter species are considered as a serious multidrug resistant pathogen and is encompassed in the “ESKAPE” acronym, referring to *Enterococcus faecium, Staphylococcus aureus, Klebsiella pneumoniae, Acinetobacter baumannii, Pseudomonas aerginosa* and *Enterbacter spp*^43^. Evidence for antibiotic resistant genes were identified in patient 6 within *Klebsiella pneumoniae* (TEM-4) and *Acinetobacter baumannii* species (LpxC, adeI, adeJ, mexT, adeN, adeK, ADC-2) (Table 5). This patient died from a severe MERS infection; the impact of the potential co-infection is not known. Several groups have suggested that the widespread of antibiotics as part of the management of patients with COVUD-19 may effect antimicrobial resistance (AMR)^44^. Metagenomic analysis of samples from patients with severe coronavirus infection, as illustrated in this study, can be used to track these markers. *Human alphaherpesvirus 1* transcripts were identified in patient 5 through this approach. Ongoing research throughout the SARS-CoV-2 pandemic suggests that coronaviruses may be able to reactivate herpes simplex virus and cytomegalovirus during severe disease^44^.

**Table 5:**
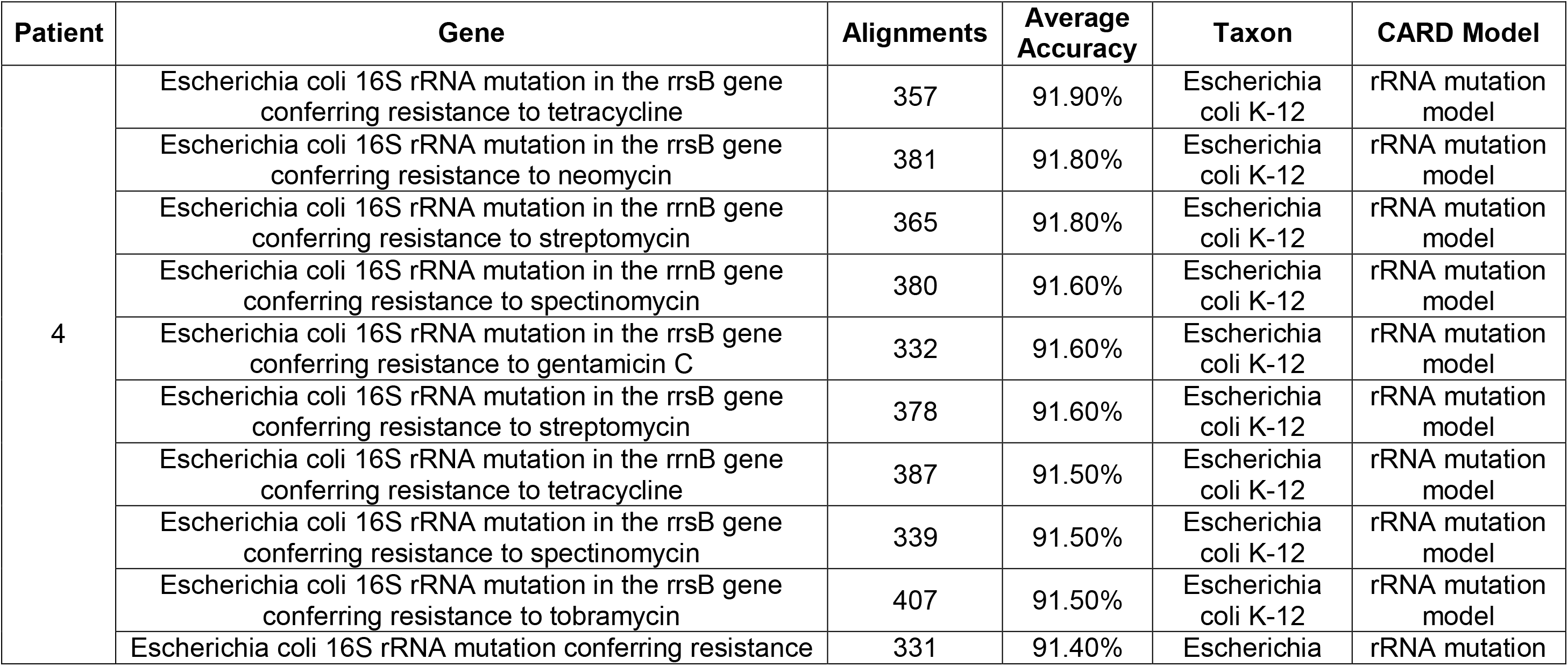
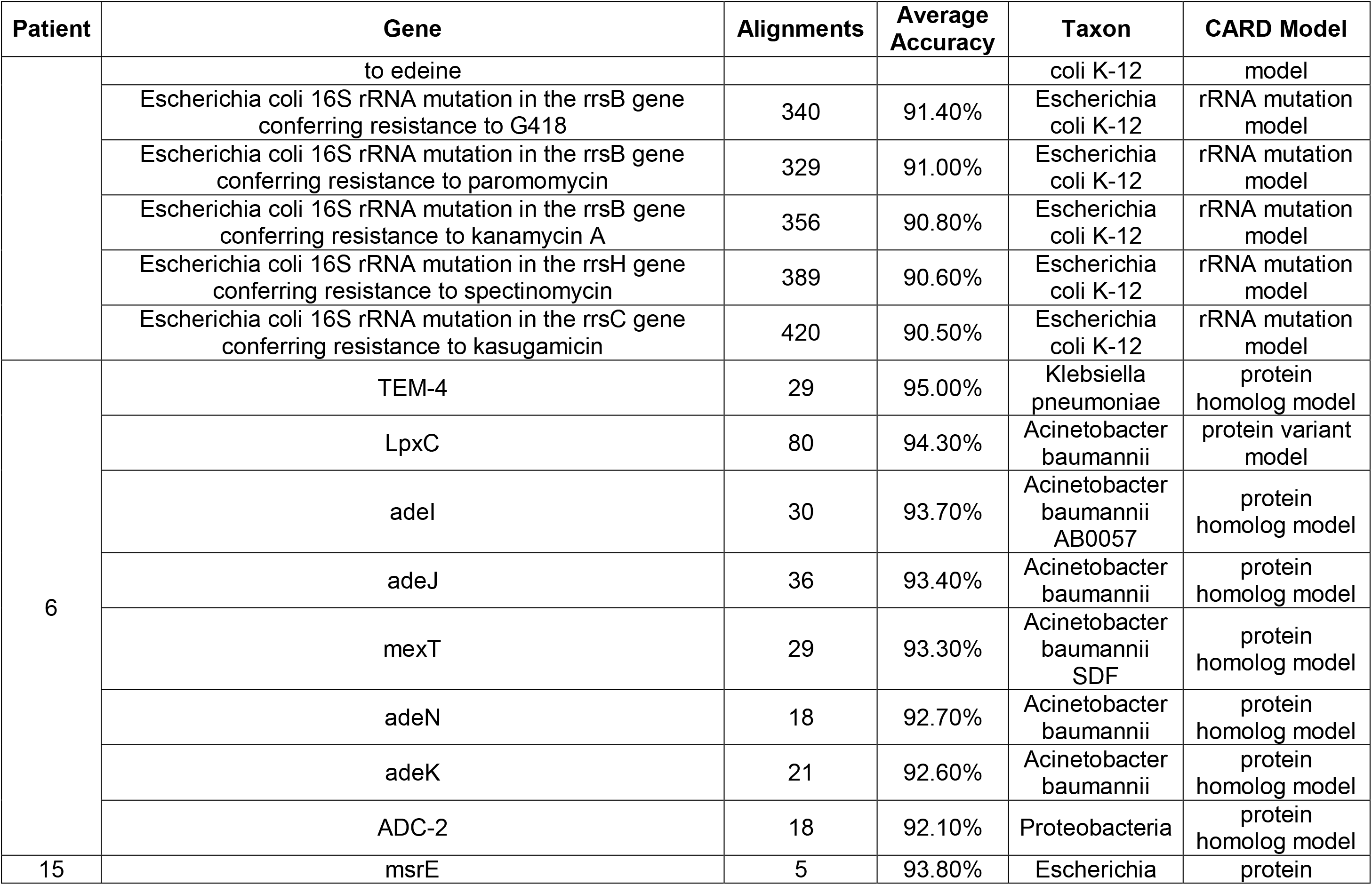
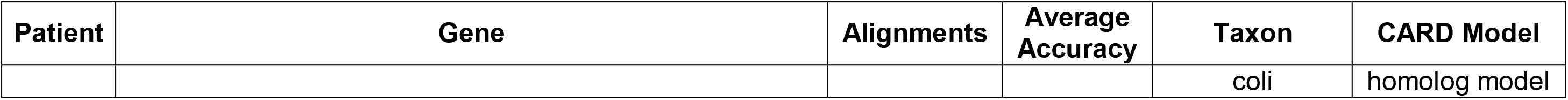
Antimicrobial resistant genes identified in bacteria from patients with MERS-CoV infection. A Sequence-Independent, Single Primer Amplification (SISPA) method was used to identify viral and bacterial transcripts within clinical samples. Fastq files were uploaded to Oxford Nanopore Technology’s (ONT’s) cloud-based pipeline EPI2ME (Fastq Antimicrobial Resistance; WIMP (rev. 3.4.0), ARMA CARD (rev. 1.1.6)) workflow to retrieve taxonomy classification and antimicrobial resistance information from MERS-CoV clinical samples. Antimicrobial resistant gene candidates with more than 5 alignments and an average accuracy of more than 90% are presented.

The abundance of bacteria was compared between fatal cases and non-fatal cases using DESeq2 (Fig. 10). In general, bacteria from the Proteobacteria phyla were in higher abundance in the fatal cases in comparison to non-fatal, including species from the Acinetobacter and Klebsiella genera. Previous studies have suggested interactions between viral infections and bacterial communities. An increase in abundance of Proteobacteria was observed following an experimental rhinovirus infection in COPD patients^45^. Patients with 2009 influenza A H1N1 infections and pneumonia, were found to have an expansion of Proteobacteria in comparison to non-viral pneumonias^46-48^. The data indicated that an amplicon-based approach could be used to rapidly sequence MERS-CoV from clinical samples and provide information on genetic diversity and insertions/deletions. This was complemented with a metagenomic approach that was able to resolve the microbiome present in the clinical sample. This analysis identified bacteria associated with ventilation and also antibiotic resistance markers. Overall, the research demonstrates the utility of rapid long read length sequence to characterise MERS-CoV infection in samples from humans with MERS.

**Fig. 10:**
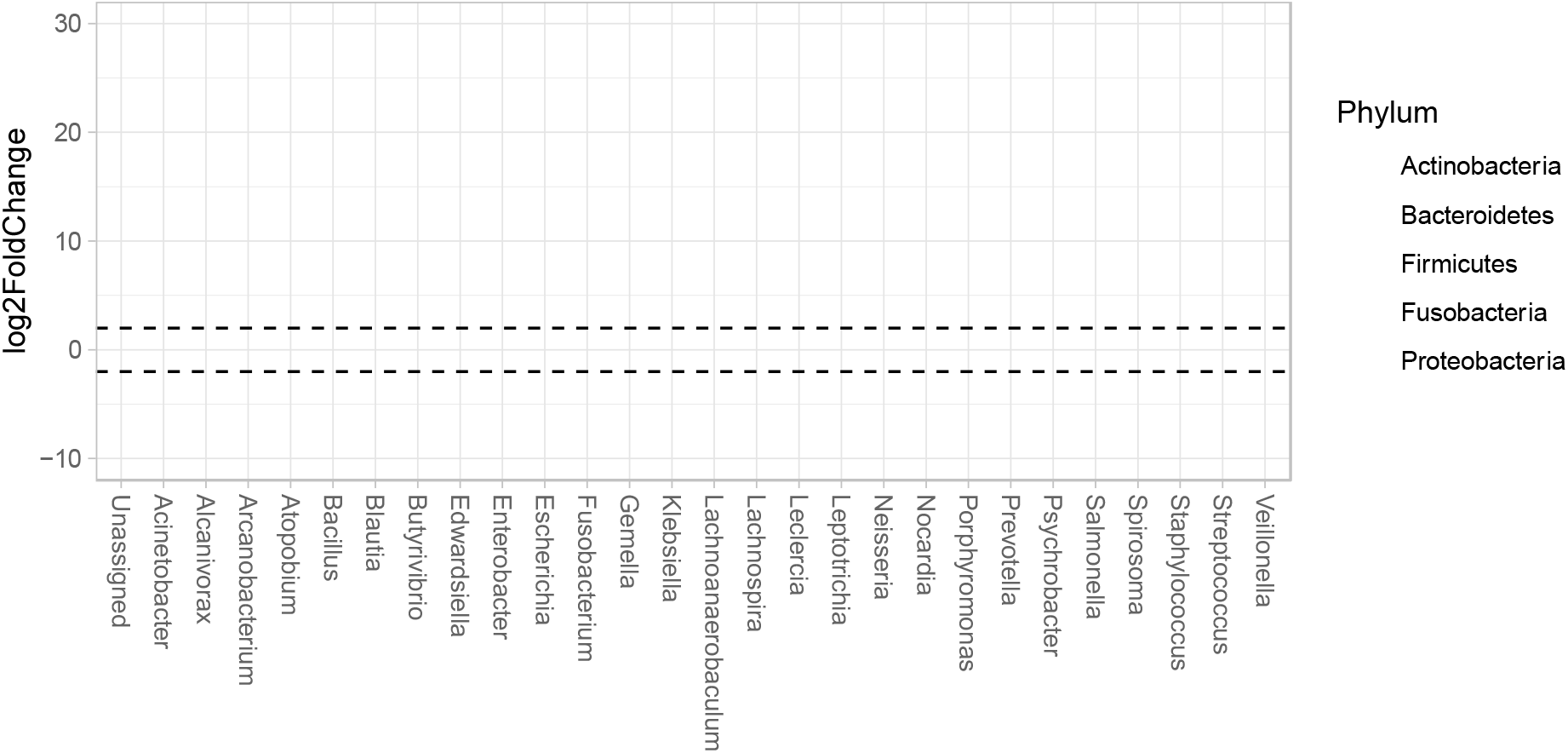
Up and down regulated abundance of bacteria in fatal cases of MERS-CoV infections in comparison to non-fatal cases. Phyloseq object was converted into a DESeq2 object and a contrast between fatal and non-fatal outcome was used to calculate log2 fold change, values with an FDR <0.01 were plotted. X axis represents the genus of identified transcripts, unassigned refers to transcripts that cannot be assigned at the genus level, however, colour illustrates the phyla.

## Acknowledgments

We would like to acknowledge the Research Center at King Fahad Medical City for funding this study (Grant no. 019-003).

## Author contributions

Conceptualisation: W.A. and J.A.H. Methodology: D.A.M., M.A., Z.M., A.D.D., W.A., R.P.-R., A.A., A.N., S.A, H.A., M.A., M.W.C. and J.A.H. Software: R.P.-R. and X.D. Validation: M.A., R.P.-R., A.J.H., Y.R. and X.D. Formal analysis: R.P.-R., D.A.M., A.D., Y.R. and X.D. Investigation: W.A., A.D.D., M.A., R.P.-R., J.A.H., I.D.-B., Y.R., Z.M., A.A., N.J.R, X.D., A.J.H., B.Y.A., D.A., B.Y.A., B.A., M.H., Resources: W.A., A.D.D., D.A.M. and X.D. Metadata: A.A., A.R.A., Data curation: R.P.-R. and X.D. Writing – Original Draft Preparation: W.A. and J.A.H. Writing – Review and Editing: all authors. Visualisation: M.A., R.P.-R., Y.R. and A.J.H. Supervision: W.A., A.D., M.W.C. and J.A.H. Project administration: W.A. and J.A.H. Funding: W.A., M.W.C. and J.A.H.

## References

1. Channappanavar R, Perlman S. Pathogenic human coronavirus infections: causes and consequences of cytokine storm and immunopathology. Semin Immunopathol 2017; 39(5): 529–39.

2. Alosaimi B, Hamed ME, Naeem A, et al. MERS-CoV infection is associated with downregulation of genes encoding Th1 and Th2 cytokines/chemokines and elevated inflammatory innate immune response in the lower respiratory tract. Cytokine 2020; 126: 154895.

3. Salmon-Rousseau A, Piednoir E, Cattoir V, de La Blanchardiere A. Hajj-associated infections. Med Mal Infect 2016; 46(7): 346–54.

4. Falzarano D, de Wit E, Rasmussen AL, et al. Treatment with interferon-alpha2b and ribavirin improves outcome in MERS-CoV-infected rhesus macaques. Nat Med 2013; 19(10): 1313–7.

5. Modjarrad K, Roberts CC, Mills KT, et al. Safety and immunogenicity of an anti-Middle East respiratory syndrome coronavirus DNA vaccine: a phase 1, open-label, single-arm, dose-escalation trial. Lancet Infect Dis 2019; 19(9): 1013–22.

6. Adney DR, Wang L, van Doremalen N, et al. Efficacy of an Adjuvanted Middle East Respiratory Syndrome Coronavirus Spike Protein Vaccine in Dromedary Camels and Alpacas. Viruses 2019; 11(3).

7. Alharbi NK, Qasim I, Almasoud A, et al. Humoral Immunogenicity and Efficacy of a Single Dose of ChAdOx1 MERS Vaccine Candidate in Dromedary Camels. Sci Rep 2019; 9(1): 16292.

8. Alharbi NK, Padron-Regalado E, Thompson CP, et al. ChAdOx1 and MVA based vaccine candidates against MERS-CoV elicit neutralising antibodies and cellular immune responses in mice. Vaccine 2017; 35(30): 3780–8.

9. Ooi PL, Lim S, Chew SK. Use of quarantine in the control of SARS in Singapore. Am J Infect Control 2005; 33(5): 252–7.

10. Ruan YJ, Wei CL, Ee AL, et al. Comparative full-length genome sequence analysis of 14 SARS coronavirus isolates and common mutations associated with putative origins of infection. Lancet 2003; 361(9371): 1779–85.

11. Liu J, Lim SL, Ruan Y, et al. SARS transmission pattern in Singapore reassessed by viral sequence variation analysis. PLoS Med 2005; 2(2): e43.

12. Carroll MW, Matthews DA, Hiscox JA, et al. Temporal and spatial analysis of the 2014-2015 Ebola virus outbreak in West Africa. Nature 2015; 524(7563): 97–101.

13. Quick J, Loman NJ, Duraffour S, et al. Real-time, portable genome sequencing for Ebola surveillance. Nature 2016; 530(7589): 228–32.

14. Dudas G, Carvalho LM, Bedford T, et al. Virus genomes reveal factors that spread and sustained the Ebola epidemic. Nature 2017; 544(7650): 309–15.

15. Diallo B, Sissoko D, Loman NJ, et al. Resurgence of Ebola Virus Disease in Guinea Linked to a Survivor With Virus Persistence in Seminal Fluid for More Than 500 Days. Clin Infect Dis 2016; 63(10): 1353–6.

16. Kafetzopoulou LE, Pullan ST, Lemey P, et al. Metagenomic sequencing at the epicenter of the Nigeria 2018 Lassa fever outbreak. Science 2019; 363(6422): 74–7.

17. Yozwiak NL, Schaffner SF, Sabeti PC. Data sharing: Make outbreak research open access. Nature 2015; 518(7540): 477–9.

18. Van Kerkhove MD, Alaswad S, Assiri A, et al. Transmissibility of MERS-CoV Infection in Closed Setting, Riyadh, Saudi Arabia, 2015. Emerg Infect Dis 2019; 25(10): 1802–9.

19. Smith EC, Case JB, Blanc H, et al. Mutations in coronavirus nonstructural protein 10 decrease virus replication fidelity. J Virol 2015; 89(12): 6418–26.

20. Subissi L, Posthuma CC, Collet A, et al. One severe acute respiratory syndrome coronavirus protein complex integrates processive RNA polymerase and exonuclease activities. Proc Natl Acad Sci U S A 2014; 111(37): E3900–9.

21. Kottier SA, Cavanagh D, Britton P. Experimental evidence of recombination in coronavirus infectious bronchitis virus. Virology 1995; 213(2): 569–80.

22. Kusters JG, Jager EJ, Niesters HG, van der Zeijst BA. Sequence evidence for RNA recombination in field isolates of avian coronavirus infectious bronchitis virus. Vaccine 1990; 8(6): 605–8.

23. Jackwood MW, Boynton TO, Hilt DA, et al. Emergence of a group 3 coronavirus through recombination. Virology 2010; 398(1): 98–108.

24. Herrewegh AA, Smeenk I, Horzinek MC, Rottier PJ, de Groot RJ. Feline coronavirus type II strains 79-1683 and 79-1146 originate from a double recombination between feline coronavirus type I and canine coronavirus. J Virol 1998; 72(5): 4508–14.

25. Sohrab SS, Azhar EI. Genetic diversity of MERS-CoV spike protein gene in Saudi Arabia. J Infect Public Health 2019.

26. Li YT, Chen TC, Lin SY, et al. Emerging lethal infectious bronchitis coronavirus variants with multiorgan tropism. Transbound Emerg Dis 2020; 67(2): 884–93.

